# Evaluating Impacts of Syntenic Block Detection Strategies on Rearrangement Phylogeny Using M. tuberculosis Isolates

**DOI:** 10.1101/2022.02.18.481113

**Authors:** Afif Elghraoui, Siavash Mirarab, Krister M. Swenson, Faramarz Valafar

## Abstract

Phylogenetic inference based on genomic structural variations, that manipulate the gene order and content of whole chromosomes, promises to inform a more comprehensive understanding of evolution. The first challenge in using such data, the incompleteness of available *de novo* assemblies, is easing as long read technologies enable (near-)complete genome assembly, but methodological challenges remain. To obtain the input to rearrangement-based inference methods, we need to detect syntenic blocks of orthologous sequences, a task that can be accomplished in many ways, none of which are obviously preferable. In this paper, we use 94 reference quality genomes of primarily *Mycobacterium tuberculosis* (Mtb) isolates as a benchmark to evaluate these methods. The clonal nature of Mtb evolution, the manageable genome sizes, along with substantial levels of structural variation make this an ideal benchmarking dataset. We test several methods for detecting homology and obtaining syntenic blocks, and two methods for inferring phylogenies, comparing them to the standard method that uses substitutions for inferring the tree. We find that not only the choice of methods but also their parameters can impact results, especially among branches with lower support. In particular, a method based on an encoding of adjacencies applied to Cactus-defined blocks was fully compatible with the highly supported branches of the substitution-based tree. Thus, we were able to *combine* the two trees to obtain a supertree with high resolution utilizing both SNPs and rearrangements. Furthermore, we observed that the results were much less affected by the choice of the tree inference method than by the method used to determine the underlying syntenic blocks. Overall, our results indicate that accurate trees can be inferred using genome rearrangements, but the choice of the methods for inferring the homology matters and requires care.

## 1 Introduction

Methods for phylogenetic inference based on genomic rearrangements have been developed and refined over the past several decades [Moret et al., 2013], but the majority of biologists continue to rely primarily on phylogenetic inference methods based on nucleotide substitutions. As a major evolutionary mechanism, genomic structural variation cannot be neglected, yet application of gene-order methods have been limited to specific cases of well-assembled eukaryotic genomes [e.g., Pevzner and Tesler, 2003, Feng et al., 2017], as well as to small plastid genomes [Moret et al., 2002]. For example, the tool CREx is currently the standard software for analyzing plastid gene order evolution [Bernt et al., 2007]. The widespread application of these methods to larger genomes, however, has faced many challenges. Before the recent advent of accurate long-read sequencing technology, the proper *de novo* assembly of full genomes for use as input to a gene order analysis had proven difficult; indeed the mechanisms of rearrangement are often inextricably linked to duplicated regions that are well known for confusing short-read reference-based sequencing methods [Ranz et al., 2007, Liu et al., 2012].

Assembly of reference-quality bacterial genomes is now routine [Koren and Phillippy, 2015, Phillippy, 2017] and full assembly of the much larger and complex eukaryotic genomes is now possible [Liao et al., 2021, Miga et al., 2019, Nurk et al., 2021, Rhie et al., 2021]. The improved assemblies have the potential to enable a wider use of rearrangements as a phylogenetic signal. However, methodological challenges associated with the use of such signal remain and call for better methods and better empirical understanding of the strengths and weaknesses of existing methods.

Much of the research in rearrangement phylogenetics is focused on the difficult problem of modeling complex scenarios that can arise from genomic structural variations mediated by many different mechanisms [Fertin, 2009]. Methods that infer phylogenies based on rearrangements are of three varieties [Moret et al., 2013]: a) Model-free methods, which treat rearrangements as character evolution. For example, some methods encode adjacencies as simple characters (e.g., binary or copy-number) and use standard character evolution models together with maximum likelihood inference [e.g., Wang et al., 2002, Lin et al., 2012a, Hu et al., 2014]. b) Another family of methods uses distance-based tree reconstruction by finding the minimum number of events needed to transform one genome to another [e.g., Moret et al., 2003, Wang et al., 2006, Feijao and Meidanis, 2011, Bohnenkämper et al., 2020], a problem that remains challenging if there are duplicated genes [Fertin, 2009]. c) Finally, models of pairwise comparison can be generalized to the computation of rearrangements on a phylogeny. Instead of solving the small phylogeny problem directly, these approaches are usually based on the median of three genomes [Sankoff and Blanchette, 1998, Pe’er and Shamir, 1998, Moret et al., 2013], sometimes coupled with other heuristics [Bourque and Pevzner, 2002]. While the last approach is the most thorough, and a version of the median problem can now often be solved for duplication-free scenarios of reasonable size [Xu, 2010], the small phylogeny problem has proved very challenging in the presence of segmental duplication. Thus, unless compromises are made that resolve duplicated segments beforehand, the first two approaches are the only types of method that are practical for datasets of even moderate size. Several algorithms exist from both categories, and the relative accuracy of these methods has been the subject of study [e.g., Lin et al., 2011, Biller et al., 2016]. Regardless of which one is used, a more prosaic question is that of preparing the input to these methods.

Although there are exceptions [Doerr et al., 2018], the input to rearrangement phylogeny reconstruction consists of a set of homologous *blocks* of nucleotide sequence, their homology assignments within and across genomes, and their relative positions and directions along the genomes. Clearly, there are many ways to define such blocks [Ghiurcuta and Moret, 2014], and detection of homology is far from trivial. The most obvious approach to homology detection is to annotate genes using gene models. This approach has to contend with difficulties of gene annotation [Salzberg, 2019], and outside prokaryotes, the more damaging issue that only a small portion of the genome can be used. An alternative is to pairwise align genomes with respect to each other or a reference genome and use the alignments to define blocks of homologous sequence. Any definition of a block should allow some levels of heterogeneity within the block, often necessitating thresholds for defining how much variability is tolerated. Pairwise alignment has a fundamental limitation: there is no guarantee that their results are consistent (e.g., are transitive). Thus, an arguably better approach is to rely on whole genome alignment (WGA). There has been much progress in recent years on scalable and accurate methods for WGA [Earl et al., 2014, Armstrong et al., 2019] and new ways to compute syntenic blocks [Kolmogorov et al., 2018].

Defining the block-level input to rearrangement phylogeny algorithms remains a challenging problem [Lucas and Roest Crollius, 2017], as demonstrated by a thorough literature search which reveals roughly thirty tools developed for that purpose since 2004. Ghiurcuta and Moret [2014] attempted to set basic standards for defining syntenic blocks but also acknowledged that the variety of criteria for selecting them corresponds to the variety of applications for which they are used, and so what is better for one application does not necessarily suit another. Despite these attempts, little is known about the relative accuracy of available options and the extent of their impact on the resulting phylogeny. One challenge when studying block inference methods is the lack of a sufficiently realistic genome simulators; to our knowledge existing simulators either do not allow for ancestral genomes to be specified as part of the input [Hindré et al., 2012, Davín et al., 2020], or are no longer maintained [Edgar et al.]. In the absence of such simulations, we have to rely on empirical data, which poses its own challenges. In particular, studying the impact of block definition would be further complicated for large genome sizes, or very complex evolutionary histories including horizontal transfers, gene duplication events, or polyploidy. Thus, a relatively simple model organism is preferable.

As a way to minimize the issue of the evolutionary model complexity, we consider *Mycobacterium tuberculosis* (*Mtb*) as a subject. Mycobacteria are unique in that they do not undergo horizontal gene transfer (HGT) in the traditional sense. While some mycobacteria have been observed to recombine via distributive conjugal transfer (DCT) [Gray and Derbyshire, 2018], the human-adapted pathogen *Mtb* in particular appears to have recently diverged yet contains appreciable diversity [Galagan, 2014, Coscolla and Gagneux, 2014a]. It does not show evidence of either DCT or traditional HGT and appears to have undergone strictly vertical evolution [Brosch et al., 2002, Gagneux, 2018]. Nevertheless, structural variations do happen for these strains; even within the species, gene duplication and gene conversion has been observed [Uplekar et al., 2011], as well as inversions [Merrikh and Merrikh, 2018]. According to the classification of Koren et al. [2013], *Mtb* has a class II genome, characterized by many mid-scale repeats of approximately 1.5 kb insertion sequences. Focusing on *Mtb* simplifies the evolutionary models we must consider, and leaves us with the final difficulty of rearrangement phylogeny: defining suitable synteny blocks. Despite its apparent clonal evolution, *Mtb* has diversified into several lineages (Fig. 1) distinguished by variations in repetitive regions [Kanduma et al., 2003], with the three “modern” lineages further separated from four “ancestral” lineages by the deletion of the TbD1 locus [Gordon et al., 1999, Brosch et al., 2002, Mostowy et al., 2002]. Its evolution is driven in part by antibiotic pressure, though some lineages are more virulent and more successful than others [Merker et al., 2015], such as the globally prevalent Euro-American (L4) and East-Asian (L2) lineages [Coscolla and Gagneux, 2014b].

**Figure 1:**
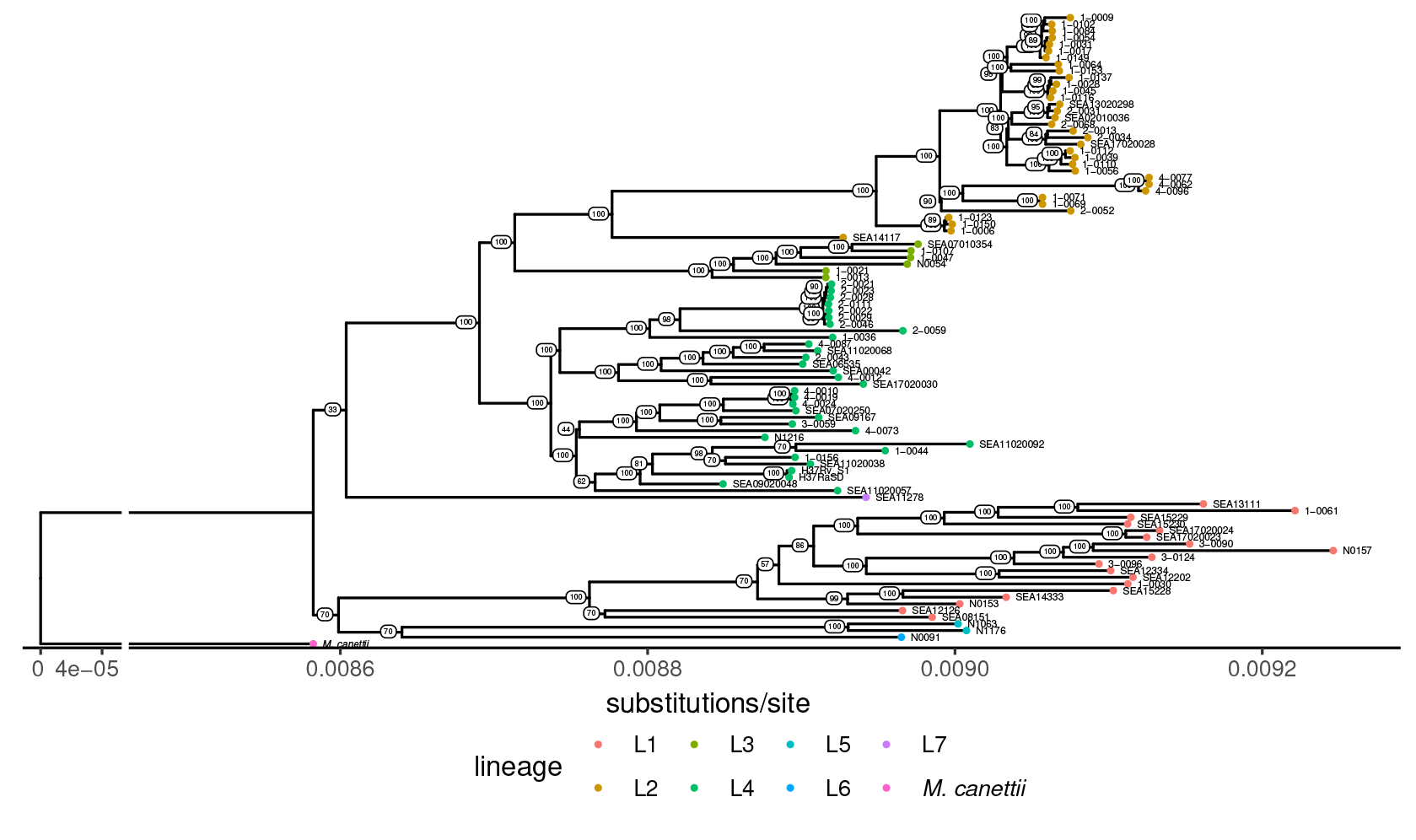
Phylogeny of *Mycobacterium tuberculosis* clinical isolates used in this study. This maximum-likelihood phylogenetic tree inferred using the GTRCAT substitutions model from their SNPs with respect to the reference strain H37Rv (NC_000962.3) shows the separation of our *Mtb* isolate set into 7 of the defined lineages and the level of diversity between them. This includes 3 *M. africanum* isolates (lineages 5 and 6). *M. canettii* is used as the outgroup.

In this article we use a set of complete genomes for 92 largely drug-resistant *Mtb* clinical isolates and two reference strains, evaluating different methods of syntenic block determination with respect to how an adjacency phylogeny built from them compares to standard trees inferred from substitutions. For each method, we use both an adjacency-based algorithm, MLWD [Lin et al., 2012a], and a recent distance-based method called DING [Bohnenkämper et al., 2020], to infer rearrangement-based phylogenies. We use two distinct approaches as input to these methods: 1) synteny blocks determined using modern WGA methods (Cactus [Armstrong et al., 2020] with different parameters and SibeliaZ-LCB [Minkin and Medvedev, 2020]), and 2) blocks determined by our in-house gene annotation pipeline. We compare the resulting trees to each other and to those inferred using substitutions alone in order to quantify their levels of discordance, especially with respect to branches with high statistical support.

## 2 Methods

### 2.1 Genome Assemblies

Genome assemblies made available by Modlin et al. [2020] are used. These include 85 isolates resequenced from the set collected by the Global Consortium for Drug-resistant Tuberculosis Diagnostics (GCDD) [Hillery et al., 2014] (NCBI Bioproject PRJNA555636), 7 isolates from Berney et al. [2015] (NCBI Bioproject PRJEB8783), reference strains H37Rv (NCBI Bioproject PRJNA555636), H37Ra (NCBI accession NZ_CP016972.1), and *Mycobacterium canettii* (NCBI accession NC_019951.1). All sequencing data used for the assemblies, except the *M. canettii* assembly, are Pacific Biosciences SMRT sequencing reads. The assembly protocol of Modlin et al. [2020] was based on HGAP2 [Chin et al., 2013] or, if that failed to produce a single contig, Canu [Koren et al., 2017]. The contigs were circularized using minimus2 [amo] or circlator [Hunt et al., 2015], followed by iterative assembly consensus polishing with Quiver [Chin et al., 2013]. To validate our assemblies, we used structural variant detection method PBHoney [English et al., 2014], which detects irregularities such as soft-clippings in the alignment of reads to a reference. We applied PBHoney to the reads’ alignment to the assembly that was generated from them, so any structural variant detected by PBHoney would in fact be a candidate misassembly. None of the genomes used here had any misassemblies detected.

Lineages were identified using TB-profiler[Phelan et al., 2019] version 4.1.1 with database version 2022-01-25.

### 2.2 Block Assignment

The synteny block assignment strategies we used here fall into the two categories of annotation-based and alignment-based methods and so differ in the type of markers they use for defining their blocks.

#### 2.2.1 Annotation-Based Methods

##### Annotation and Homology Assignment

All genomes were simultaneously annotated using the Hybran pipeline [hyb]. Hybran uses a combination of reference-based annotation transfer implemented by RATT [Otto et al., 2011] and Prokka, an *ab initio* method [Seemann, 2014]. Annotations from Prokka are only retained in the places of gaps left where no suitable reference annotation could be transferred. The reference annotation used was that of *M. tuberculosis* H37Rv (NCBI accession NC_000962.3).

In the last stage of the annotation pipeline, orthologous genes across the genomes are identified using CD-HIT [Li and Godzik, 2006, Fu et al., 2012] and MCL [Enright et al., 2002] clustering. Genes that cluster together are assigned the same name if they have at least 95% protein sequence identity and 95% alignment coverage with the representative sequence of the cluster (this threshold is applied to both query/subject and subject/query coverage). These arbitrary thresholds clearly have the potential to impact results. Thus, we also ran the annotation pipeline with relaxed thresholds for the orthology mapping: 75% minimum identity and 66% minimum alignment coverage. These results are subsequently referred to as “annotation-relaxed”.

### 2.2.2 Alignment-Based Methods

#### Cactus

[Armstrong et al., 2020] is a whole-genome aligner based on its namesake cactus graphs, which organize alignments hierarchically to reveal their substructures. The Cactus aligner is run with default parameters. Cactus requires soft-masking of repetitive sequences prior to alignment, so we applied nucmer [Kurtz et al., 2004] to identify them and bedtools [Quinlan and Hall, 2010] to apply the masking. Cactus requires a guide tree as input, and the reference SNP tree (described in Section 2.4) was provided by default. We tested the robustness to this choice by changing the guide tree. We constructed an alternative guide tree using FastME [Lefort et al., 2015] on a distance matrix computed using Mash [Ondov et al., 2016a] using the maximum k-mer size (-k) of 32 due to the clonality of the dataset, and a sketch size (-s) of 1 billion, and phylogenetically corrected using the Jukes and Cantor [1969] (JC) model. The guide tree used is denoted in parentheses if it is not the SNP tree. For example, Cactus(Mash) refers to the Cactus alignment based on the Mash guide tree.

#### SibeliaZ

[Minkin and Medvedev, 2020] is a whole-genome alignment method for closely-related genomes based on de Bruijn graph analysis. It was run with graph order (*k*-mer length) set to 15, the developer recommended value for bacterial datasets. Given that we required only the coordinates of the locally collinear blocks, rather than the full nucleotide alignments themselves, we ran SibeliaZ-LCB only, rather than the complete SibeliaZ alignment pipeline, (-n) following the developer recommendation.

#### 2.2.3 Building synteny blocks

Cactus and SibeliaZ-LCB each produce a collection of sets of alignable genomic intervals that are inferred to be homologous, called *alignment blocks*. We used a custom script (https://gitlab.com/LPCDRP/syntement/>) to formulate these alignment blocks as synteny block permutations for direct input into tree inference, as well as for formulating genes from the annotation output as such. The whole genome alignment blocks can be very short, so maf2synteny [Kolmogorov et al., 2018] was developed to process them into larger syntenic blocks. This tool aggregates multiple adjacent small alignments into larger blocks if they are consistently syntenic across genomes, and considers the lengths of the alignments and the length of the gaps between them. Specifically, maf2synteny takes two arguments: a set of simplification parameters *S* = {(*minBlock*_1_, *maxGap*_1_), (*minBlock*_2_, *maxGap*_2_), … *}* which govern when a block is expanded to incorporate adjacent aligned segments, and a synteny block scale (block_sizes), which we explore in Section 3. The method is based on an A-Bruijn graph, where there is a vertex for each alignment block (of length at least *minBlock*) and, for each genome, an edge between adjacent alignment blocks. Thus, a vertex can have maximum degree twice the number of genomes. A *collinear* path in the graph is one that includes only alignment blocks sharing the same set of genomes, and it indicates a set of collinear alignment blocks in those genomes. Maximal collinear paths, with the condition that no adjacent alignment blocks are more than *maxGap* apart, are aggregated. To permit heterogeneity within the syntenic blocks, maf2synteny also aggregates each set of alignment blocks that participate in a *bubble* (subject to the same *maxGap* parameter), which is a pair of collinear paths that share endpoints. The set of parameter pairs *S* is visited from smallest pairs to largest, and applied to the graph until the target syntenic block scale block_sizes is reached.

### 2.3 Evaluations

Because we analyze a real dataset, all trees in this study are inferred from the data. To use as a reference, we inferred trees using substitution models (Section 2.4). Clearly, there is no guarantee that these trees represent the true evolutionary history, and we invite readers to keep this point in mind when interpreting results. Nevertheless, when a method shows more similarity to the SNP tree, especially among highly supported branches, we can interpret this similarity as combined evidence for a branch in the true history. One caveats is the use of SNP tree as the guide tree for Cactus, which will be explored.

For each pair of fully resolved trees, we compare the trees using the normalized Robinson and Foulds [1981] (RF) distance and matching split distance [Bogdanowicz and Giaro, 2012] (MS) metrics, computed using TreeCmp [Bogdanowicz et al., 2012]. Because we study the evolution of relatively closely related genomes, not every branch can be resolved with high confidence. Thus, in addition to comparing fully resolved trees, we also study highly supported branches. We considered each tree after contracting branches with bootstrap support (BS) below levels 0%, 33%, 50%, 75%, 95%, and 100%. For contracted trees, RF and MS become difficult to interpret. Thus, instead, we report the total number of non-trivial branches in the tree (indicating its resolution), and the number of non-trivial branches that are compatible between two trees (recall that two bipartitions are compatible if they can both exist in the same tree).

### 2.4 Phylogenetic Tree Inference

#### Reference SNP Tree

SNPs were called with respect to the reference H37Rv (NCBI accession NC_000962.3) using show-snps from the MUMmer package [Kurtz et al., 2004], then concatenated into a PHYLIP-formatted alignment. From this, RAxML [Stamatakis, 2014] was used to create a maximum-likelihood tree using the GTRCAT model with Felsenstein ascertainment bias correction using the count of sites in the reference genome where no SNP was observed, and bootstrapping with 100 replicates.

#### Rearrangement-based Trees

For each of the block assignments specified in Section 2.2, we test both an adjacency-based and distance-based method. Among adjacency-based methods, we used a tree inferred using MLGO [Hu et al., 2014] phylogenetic tree reconstruction with bootstrapping on 100 replicates. This tool implements the maximum likelihood on whole-genome data (MLWD) algorithm [Lin et al., 2012a], in which ordered markers are converted into a vector representing the copy-number of marker adjacencies. A maximum likelihoood tree is then inferred, considering the transitions between these two states for each adjacency. Among distance-based methods, we used DING [Bohnenkämper et al., 2020] to calculate a distance for every pair of samples based on the block assignments for each method (Section 2.2). DING computes distances under the double cut and join (DCJ) model that also accounts for duplications and segmental insertions/deletions. We reconstructed phylogenetic trees based on these distance matrices using FastME [Lefort et al., 2015]. The distance matrices could not be computed for the unaggregated annotation blocks or the unfiltered and unaggregated Cactus blocks after running DING for one week on a machine with 256GB with RAM, but we were able to infer distance trees for the remaining configurations: annotation+maf2synteny, Cactus-filtered and SibeliaZ-LCB (both with and without maf2synteny), which we subsequently refer to with a +DING suffix.

#### Combining trees

After contracting low support branches, we are left with multifurcating trees. When two multifurcating trees are compatible, they can be easily combined by simply finding the union of their bipartitions, which implies a tree. We built such combined trees using Dendropy [Sukumaran and Holder, 2010] and a custom script.

## 3 Results

### 3.1 Composition of syntenic blocks

While the blocks produced by all the methods cover at least 75% of the genome, the Cactus alignment resulted in complete coverage. This full coverage is achieved through the creation of an order of magnitude more syntenic units than the other methods pre-aggregation. However, Cactus coverage becomes comparable to SibeliaZ once we filtered the nearly 10,000 alignments with fewer than 50 sites (Cactus-filtered) (Table 1) that accounted for 1.1% of the genome on average. The number 50 was chosen here as it was the minimum length of blocks produced by SibeliaZ-LCB. Even still, Cactus-filtered has over double the number of blocks as SibeliaZ-LCB (7527 vs 3276).

**Table 1:** Characteristics of the Blocks Generated by each Method

| Method | Number of Blocks | Average Genome Coverage | Number of Duplicate Blocks | Number of Duplicate Occurrences | Average Multiplicity | Maximum Multiplicity |
| --- | --- | --- | --- | --- | --- | --- |
| Cactus | 17115 | 100% | 3165 | 75899 | 2.047 | 4 |
| Cactus+maf2synteny | 741 | 93.7% | 41 | 821 | 2.037 | 3 |
| Cactus-filtered | 7527 | 98.9% | 532 | 11570 | 2.020 | 3 |
| Cactus-filtered+maf2synteny | 791 | 93.6% | 41 | 821 | 2.037 | 3 |
| SibeliaZ-LCB | 3276 | 98.8% | 229 | 4217 | 2.035 | 6 |
| SibeliaZ-LCB+maf2synteny | 682 | 93.5% | 47 | 711 | 2.055 | 4 |
| annotation | 6403 | 88.6% | 35 | 3002 | 6.360 | 23 |
| annotation+maf2synteny | 2713 | 76.4% | 20 | 1441 | 7.315 | 23 |
| annotation-relaxed | 5305 | 89.0% | 50 | 3491 | 4.828 | 23 |
| annotation-relaxed+maf2synteny | 1658 | 78.6% | 26 | 1777 | 4.689 | 23 |

Gene orthology mapping using relaxed thresholds (annotation-relaxed) produced 17% fewer blocks compared to the default thresholds while maintaining the same genome coverage, reflecting the fact that more pairs of genes are identified as orthologous as a result of the lower similarity thresholds. Most blocks appeared only once in each genome (i.e., were not duplicated), in contrast to pre-aggregation Cactus and SibeliaZ, both of which have at least hundreds of duplicates. Cactus, prior to filtering, had thousands of duplicate blocks with tens of thousands of occurrences. Most of these, however, were among the short aligned segments (often only a few bp long) excluded in Cactus-filtered, and so the number of duplicate blocks drops substantially–from 3165 to 532–after filtering. The alignment-based methods did not capture into a single unit specific high-duplicity markers such as the annotated transposases, which had a maximum multiplicity of 23 in the annotations while the alignment-based methods did not identify markers with a copy number over 6 in any single genome.

With regard to running time, the three methods were widely different, with SibeliaZ-LCB taking only minutes while annotations requires hours and Cactus close to a day (Table S1). Note that Cactus is the only method here that is producing a complete WGA, explaining its increased running time.

### 3.2 Impact of block aggregation with maf2synteny

Aggregating the markers with maf2synteny generally resulted in a slight drop in coverage and, for the alignment-based methods SibeliaZ-LCB and Cactus, produced approximately 700 synteny blocks (Table 1). The coverage drop following maf2synteny for the annotation markers was more severe, from 88.6% to 76.4%, and the number of blocks was reduced by more than half (6403 to 2713). The number of duplicates also drops sharply as a result of maf2synteny, never exceeding 50 in any condition after aggregation whereas it could be as high as 3165 prior to maf2synteny. In fact, the number of duplicate markers according to the aggregated alignment-based methods becomes similar to the raw annotation despite the order of magnitude difference in the number of blocks.

The block compositions, however, are sensitive to the parameters used with maf2synteny. In particular, maf2synteny requires a block scale (`–block_sizes`), which is set by default to 5000 and which we have set to 500 to be approximately half the size of the average gene in the reference genome annotation. Exploring this parameter, we detected that it has a major impact on the number of blocks obtained, with the default resulting in 24 – 123 blocks. Reducing the block scale to 500 increased the number of syntenic blocks to around 700 for the alignment-based methods, with only minimal changes to the coverage. For all methods, the highest coverage is attained at the lowest block scale tested, 50, and the next highest coverage comes with block scale 1000 rather than 500. This is likely due to the interaction of the block scale parameter and maf2synteny’s simplification parameters (Section 2.2.3), where one more iteration of simplification takes place when using block scale 1000 versus block scale 500. The simplification parameters, however, are also tuneable to circumvent this lack of monotonicity between the block scale intervals we tested. The number of blocks and duplicate block occurrences decreases monotonically with increasing block scale for all methods. Thus, the impact of this parameter is mostly in how aggressively blocks are combined and not in how much of the genome is captured, except for annotation. Because the default value, 5000, resulted in approximately 50 blocks, it provides very little phylogenetic signal; a tree computed for Cactus using this default setting had almost no resolution with a mean BS of 8% and only four branches with BS above 60%. The coverage difference between our setting of 500 and the slight improvement seen with 1000 did not strongly justify switching to it, as the data at this point were already tractable and further simplification would result in some further loss in signal, as seen in the extreme case of 5000.

By changing the composition of the blocks, method settings also impact the final tree in substantial ways. An increased number of blocks tends to result in trees with lower branch lengths. For example, the tree inferred from relaxed annotations has less than half of the total branch length of the default annotations (0.053 vs 0.162), despite having only 20% fewer blocks. These changes are not necessarily surprising, because branch lengths are computed in the unit of mutations *per site*, which in this case can be interpreted as the number of changes *per adjacency*. When large blocks are divided into smaller blocks with little or no change in adjacencies, fewer changes will be observed per adjacency. Thus, the interpretation of branch lengths is very much tied to parameters.

### 3.3 Impact of block assignment method on trees

The choice of the block assignment methods had substantial impact on the resulting trees, especially among their less supported branches (Fig. 3). Tree resolution (i.e., the number of branches left after contracting low support branches) drops substantially at higher levels of support for rearrangement-based methods with the notable exception of Cactus-filtered without maf2synteny. The SNP tree has the highest resolution, with 93% mean BS and 63 (71) out of 91 branches having 100% (*≥* 95%) BS. Among the unaggregrated blocks, the Cactus-filtered tree has the highest resolution, followed by SibeliaZ-LCB and annotation (mean BS: 98%, 84%, and 79%, respectively). Moreover, maf2synteny reduces resolution. Considering only the trees built from maf2synteny-aggregated blocks for each method, Cactus-filtered again has the highest, followed by annotation then SibeliaZ-LCB (mean BS: 82%, 79%, and 77%, respectively).

**Figure 2:**
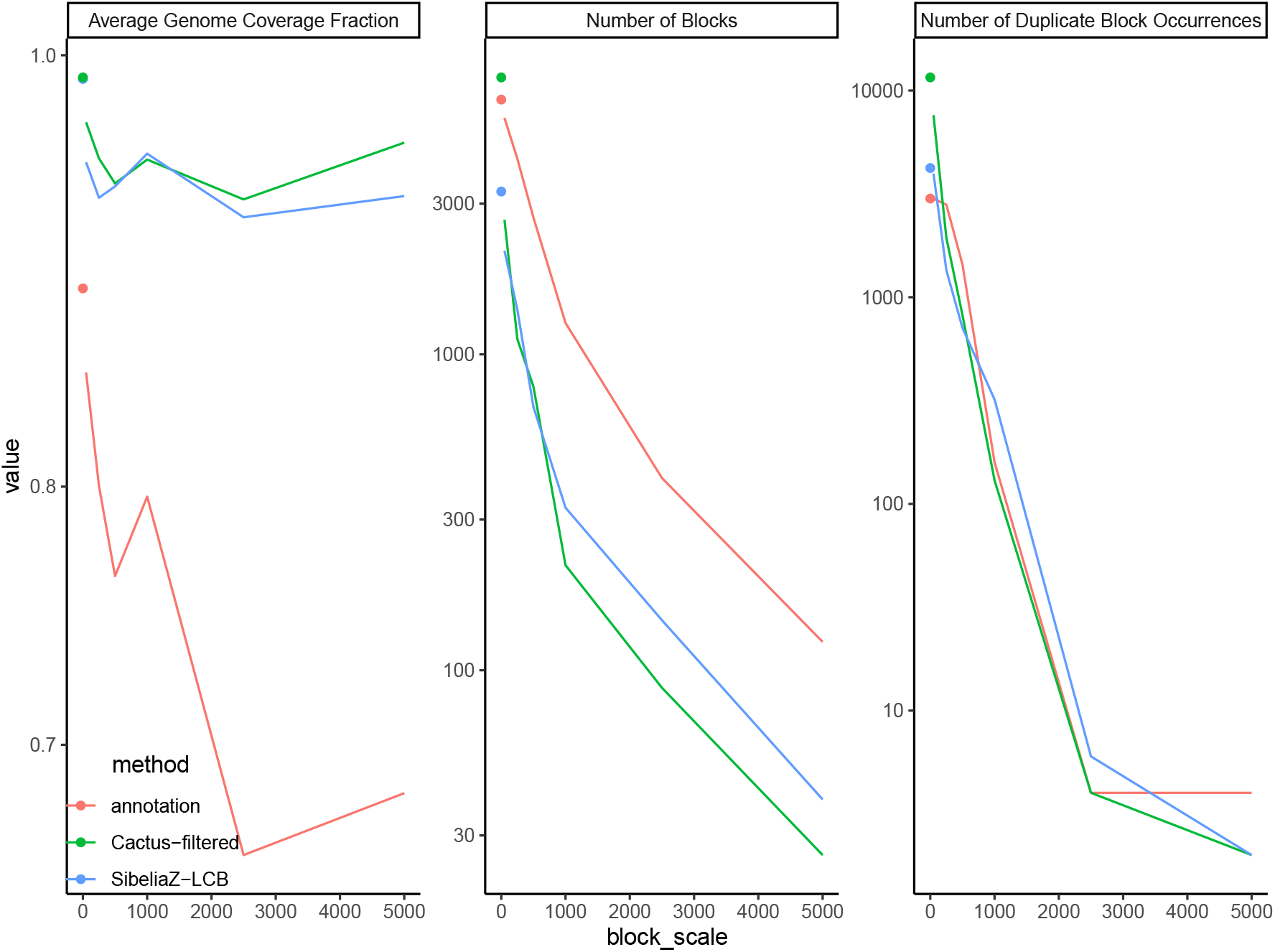
maf2synteny Parameter Effects on Synteny Blocks. Shown here are the effects of varying the `––block_sizes` parameter of maf2synteny (default 5000) on the average genome coverage fraction, resulting number of synteny blocks, and the number of duplicate occurrences. The isolated points represent the values for the raw markers. Average coverage generally decreases with increasing block scale, with the exception of a slight increase observed at block scale 1000. The overall drop in coverage following maf2synteny is more substantial for annotation and approaches 65% at larger block scales. The number of synteny blocks decreases with increasing block scale as multiple smaller blocks are combined into fewer larger blocks. Cactus and SibeliaZ-LCB have comparable numbers of blocks at each block scale The number of duplicates rapidly drops with increasing block scale for the alignment-based methods and more steadily for annotation.

**Figure 3:**
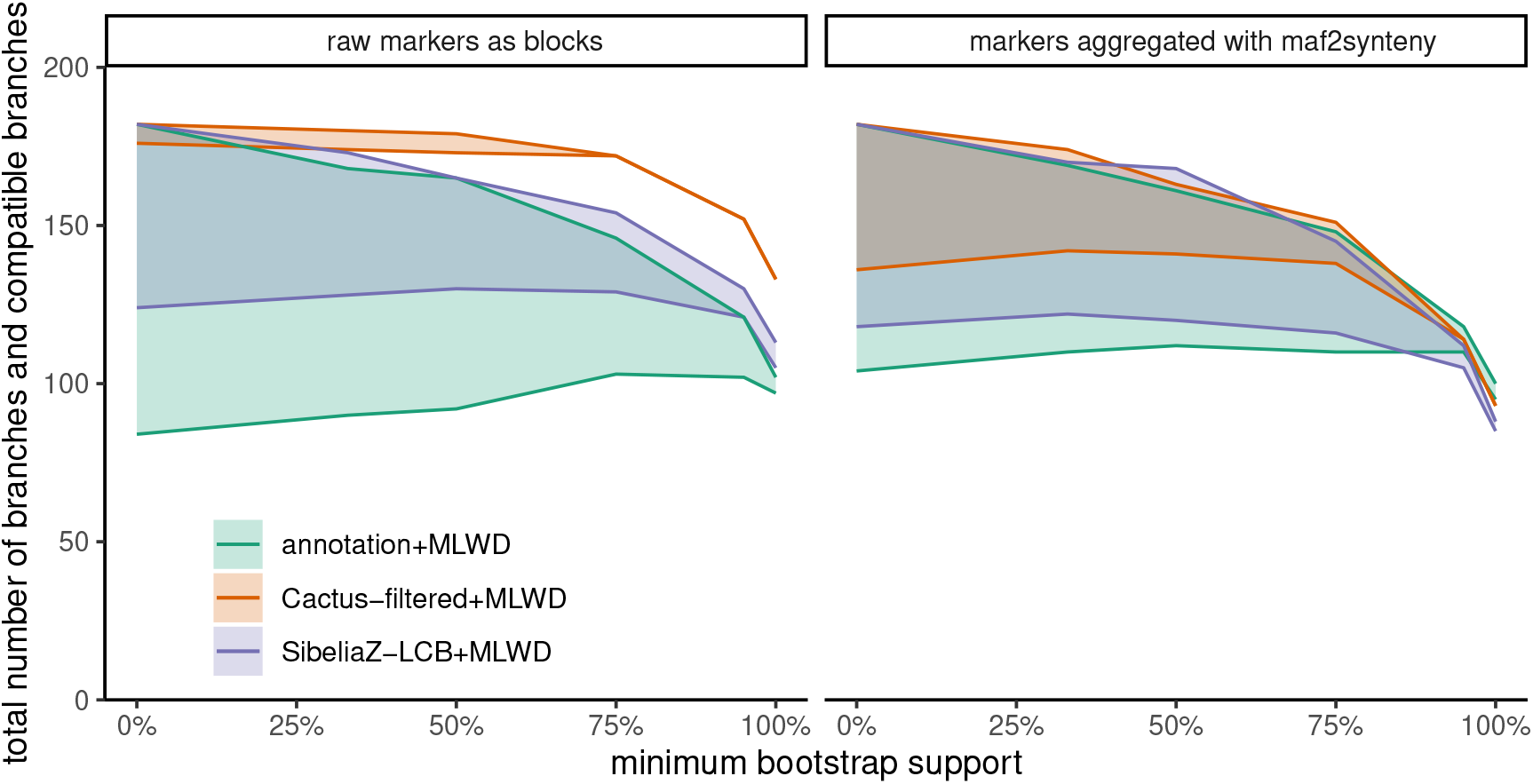
Tree Resolution and Branch Compatibility versus Minimum Bootstrap Support. Two lines are plotted for each method. The upper lines represent the total number of branches in each tree and the reference SNP tree after contracting branches with support below a threshold (*x*-axis). The lower lines represent the total number of compatible branches in the two trees. Convergence of the two lines indicates perfect compatibility. The left panel uses the MLWD trees based on the raw markers produced by the method, while the right uses markers aggregated using maf2synteny.

Note that lower support should not be interpreted as less accuracy because if a method produces incorrect synteny blocks, the resulting tree can have high support for the wrong branches. A better measure of accuracy is the compatibility of trees with the reference trees. Taking the SNP tree as reference, we observe relatively high levels of compatibility with the reference tree among adjacency-based methods and less so with annotation-based tree (Fig. 3).

Compared to Cactus-raw, Cactus-filtered had similar compatibility with the SNP tree (Fig. S2) and the added benefit of computational feasibility for running DING (the raw Cactus results in extremely large number of duplicate blocks that precluded running DING); so Cactus-filtered is exclusively used. The Cactus adjacency tree showed the greatest compatibility with the reference SNP tree, converging to perfect compatibility among branches with ≥ 75% bootstrap support. Applying maf2synteny to it, however, substantially reduces compatibility. Perfect compatibility is not achieved except among branches with *≥* 95% BS and this furthermore comes at a cost of much lower resolution.

Upon further examination of the Cactus adjacency tree (Fig. 4A), several patterns emerge. The Cactus adjacency tree is consistent with all assignments to standard lineages. Branches separating lineages are relatively long, especially for the East-Asian lineage (L2), separated by a branch of length 0.002 (i.e., 0.2% of block adjacencies have shifted in this clade). There is high BS (100%) for Indo-Oceanic (L1) to be the first to diverge from the rest, consistent with its classification as an ancestral lineage [Brosch et al., 2002, Gagneux, 2018]. There is also strong support (100%) for uniting the East-African-Indian lineage (L3) with East-Asian. The diameter of the tree is 0.01, showing that 1% of adjacencies are different between the most divergent pair of isolates. The mean distance between any pair of sequences is 0.0058.

**Figure 4:**
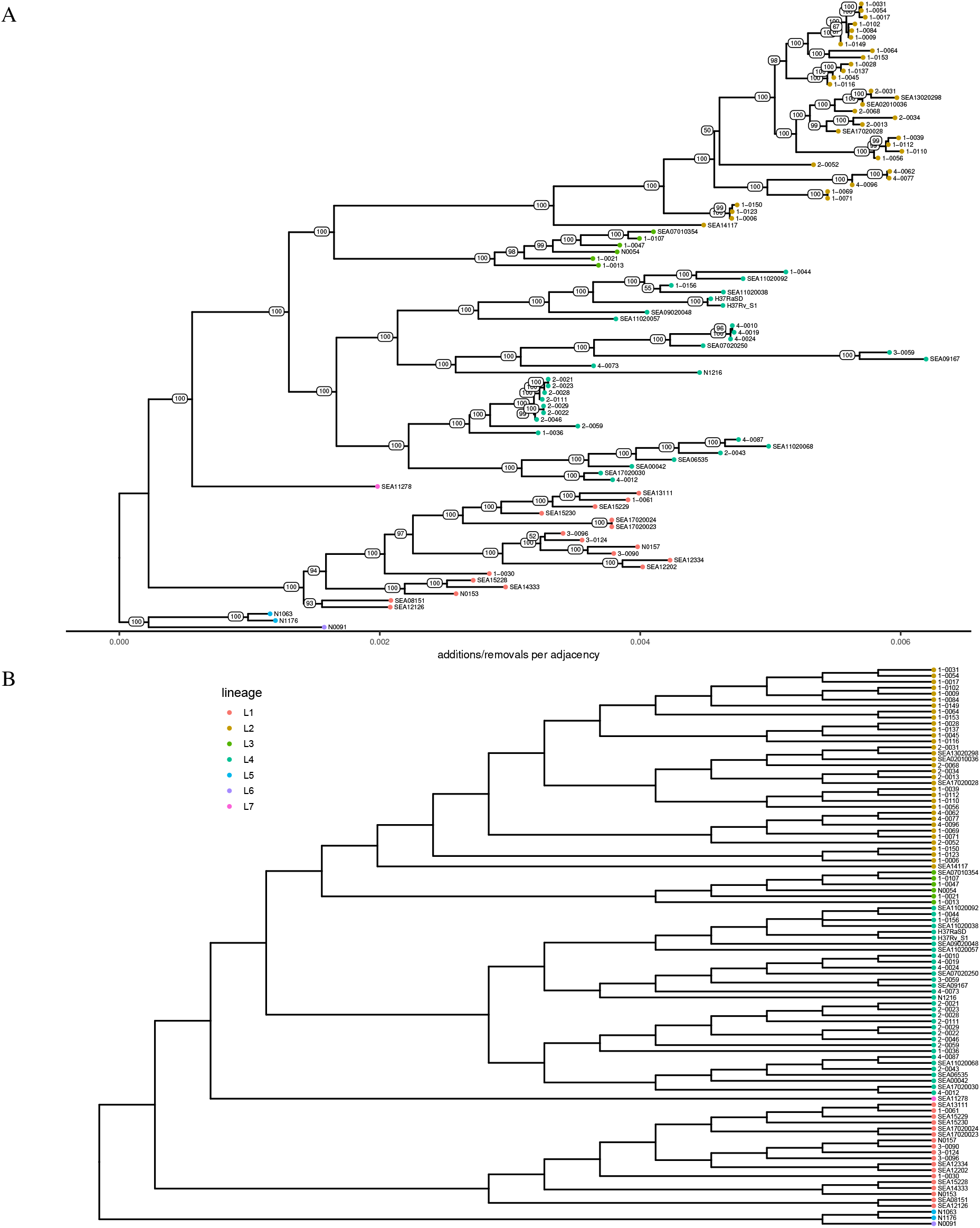
(A) Adjacency tree produced from filtered Cactus synteny blocks. (B) Combination of compatible, highly supported branches from the Cactus adjacency tree and SNP tree. Since the two trees were fully compatible for branches with ≥ 75% bootstrap support, they are easily combined to to form this tree with greater resolution. Diagrams were drawn using ggtree [Yu et al., 2017].

SibeliaZ-LCB’s adjacency tree was less compatible than Cactus with the SNP tree before aggregation. It had four (five) incompatible branches at 100% BS (*≥* 95%) with the SNP tree. maf2synteny slightly improved its compatibility, leaving one (three) incompatible branch(es) at 100% BS (*≥* 95%), but at a cost of further reducing resolution of the tree at 100% BS (32% vs 59% resolution for after and before maf2synteny, respectively). The deterioration in resolution of the alignment-based markers’ adjacency trees following maf2synteny is suggestive of reduced signal as a result of merging of orthologous alignment blocks across the taxa.

Annotation also had substantial incompatibilities with the SNP tree (nine branches at 95% BS and two branches at 100% BS), which improved with maf2synteny to four incompatible branches at 95% BS and 3 three at 100% BS, with little loss in resolution compared to the unaggregated annotation. Comparing relaxed annotation and the default annotation, we did not observe a reduction in incompatibilities with the SNP tree (Fig. S1), leading us to focus on the default annotation for the rest of the analyses. These high levels of fully supported incompatibilities in the raw annotation tree are consistent with it including strong but incorrect signal, perhaps as a result of incomplete orthology assignments resulting in missing adjacencies. The improvement in its compatibility with the SNP tree following maf2synteny may indicate that some of the incomplete and incorrect orthology assignments are masked by aggregation, making the final result a net improvement.

#### Combining complementary signals

Because the Cactus tree is fully compatible with the SNP tree at 75% BS, the two trees can easily be combined into a supertree (Fig. 4B) where every branch has *≥* 75% BS in at least one of the two source trees. This supertree makes it clear that the SNP tree and the Cactus adjacency tree include complementary signals; while the two base trees have 85 and 87 branches with *≥* 75% BS, the supertree has 90 out of 91 such branches (i.e., is 99% resolved). This supertree has all main lineages as monophyletic and the overall topology matches current understanding of Mtb’s evolution [Gagneux, 2018]: separating an East-African-Indian (L3) + East-Asian (L2) clade, first from Euro-American (L4), and then from the remaining lineages. The only remaining polytomy in the combined tree is between isolates 1-0156, SEA11020038, and the pair SEA11020092 and 1-0044. Even here, however, both the SNP and Cactus trees resolve it the same way, the former tree with BS 29% and the latter with BS 55%.

### 3.4 Impacts of guide tree

While the Cactus adjacency tree was highly compatible with the SNP tree, a caveat is that the guide tree used to infer the Cactus WGA *is* the SNP tree. Thus, to examine the impact of the Cactus guide tree on these results, we reran Cactus with two alternative guide trees: An independent one based on Mash distances (see Section 2.2.2) and the SibeliaZ-LCB+maf2synteny adjacency tree. The results show that Cactus is indeed sensitive to the guide tree. Judging by matching split (MS) distance (Fig. 5), we see that the most similar trees to each other, by a large margin of at least 50 points, are those between a Cactus adjacency/distance tree and its guide tree or between the Cactus adjacency and distance trees based on the same alignment (explored more generally in Section 3.5). While the choice of the guide tree does not impact the amount of the resolution of the Cactus adjacency tree, it dramatically impacts the topology, especially for branches with lower resolution. While Cactus(SNP)-filtered shares 97% of the SNP tree branches, Cactus(Mash)-filtered shares 81%, and Cactus(SibeliaZ-LCB)-filtered only shares 64%. Similar patterns are observed at higher bootstrap support. Among branches with *≥* 75% BS, while Cactus(SNP)-filtered had no incompatibility with the SNP tree, Cactus(SibeliaZ-LCB)-filtered has 31 such incompatible branches at *≥* 75% BS level and even 12 incompatible branches at 100% BS. Similarly, while Cactus(SibeliaZ-LCB)-filtered has only one incompatibility with the SibeliaZ-LCB guide tree at *≥* 75% BS, Cactus(SNP)-filtered has 16 incompatible branches with SibeliaZ-LCB at that level, 5 at *≥* 95% BS, and 2 at 100% BS (Fig. S3). Thus, each Cactus tree is most congruent with its own guide tree and quite incongruent with other trees, even at high support. In other words, the impact of the guide tree on the adjacency-based tree resulted in a bias towards the guide tree (as opposed to noisy variations). The fact that the impact of the guide tree extends to highly supported branches shows that the choice of the guide tree is critical.

**Figure 5:**
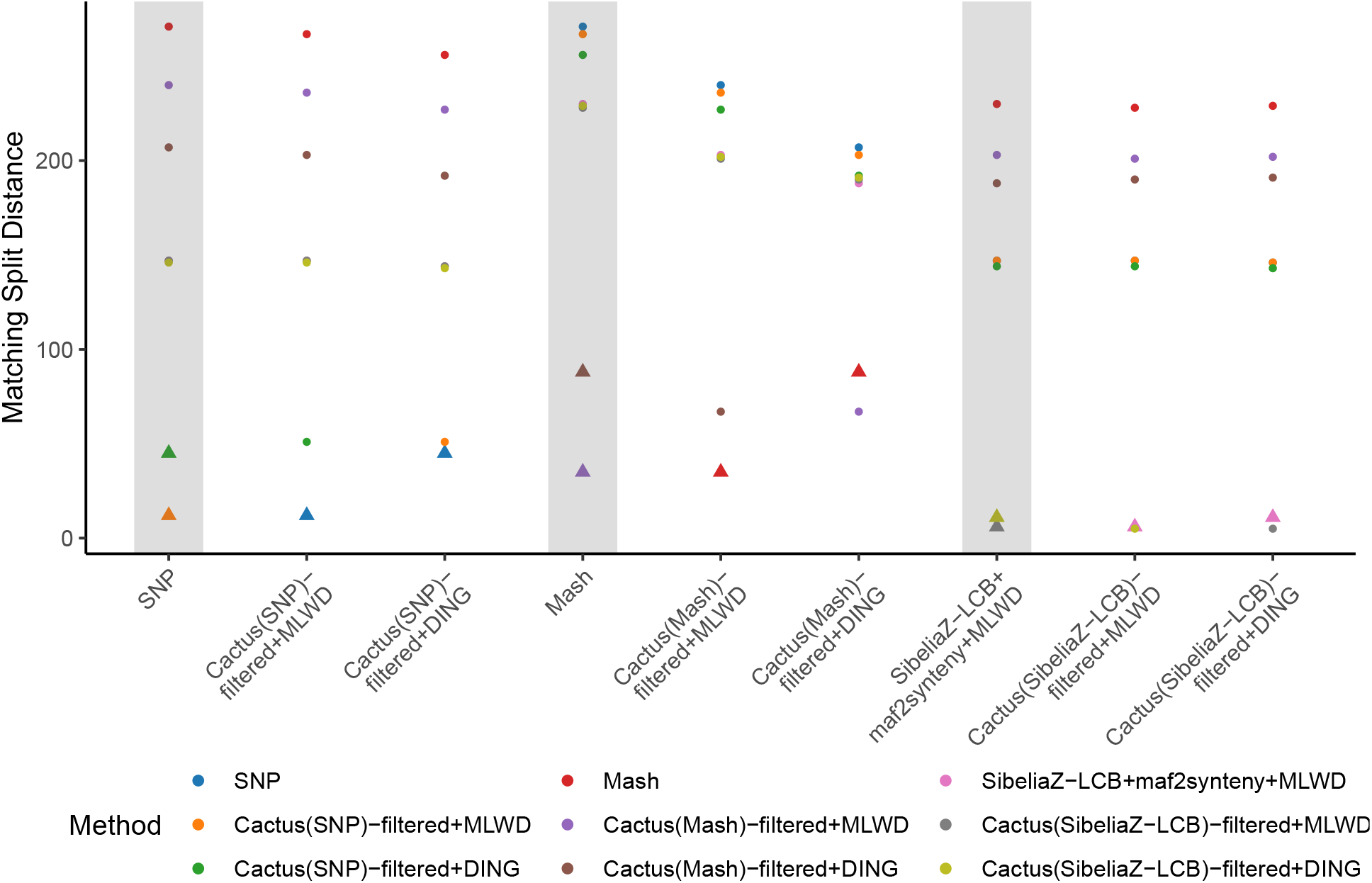
Matching split (MS) distances between trees inferred from Cactus alignments based on different guide trees. Each point corresponds to the MS distance between the method indicated on x-axis and that denoted by the point’s color. The MS distances between MLWD/DING trees and the corresponding guide tree are shown as triangular points. The most similar tree to a given MLWD/DING tree is invariably either the guide tree used for the underlying Cactus alignment itself, or the alternative tree built using the same synteny blocks (i.e. *methodA*+MLWD and *methodA*+DING). Furthermore, there is a large difference between these and the remaining independent Cactus results.

### 3.5 Impact of tree inference method

Having established that the choice of methods of block assignment do indeed matter, we next ask whether the results are robust to the choice of the tree estimation method. As bootstrapping is not commonplace for distance-based genome rearrangement methods (an approach has been proposed [Lin et al., 2012b], but not, to our knowledge, available as a software package), we compare the distance trees only to the fully resolved adjacency trees using MS and RF metrics. Overall, the distance-based and adjacency-based trees inferred from the same blocks were most similar to each other (Fig. 6), making the underlying synteny blocks the largest factor in the agreement of the trees. For example, the highest RF distance between any pair of methods was between Cactus(SNP)+maf2synteny+MLWD and annotation+MLWD methods at 56% (Fig. S4). Interestingly, the MS distances of DING trees (annotation+maf2synteny+DING, SibeliaZ-LCB+maf2synteny+DING, and Cactus+DING, both with and without maf2synteny) to the SNP tree did tend to be higher than those of the corresponding trees with MLWD. For SibeliaZ-LCB (unaggregated), the MLWD tree had a slightly higher MS distance than the corresponding DING tree (161 vs 158) (Fig. S5).

**Figure 6:**
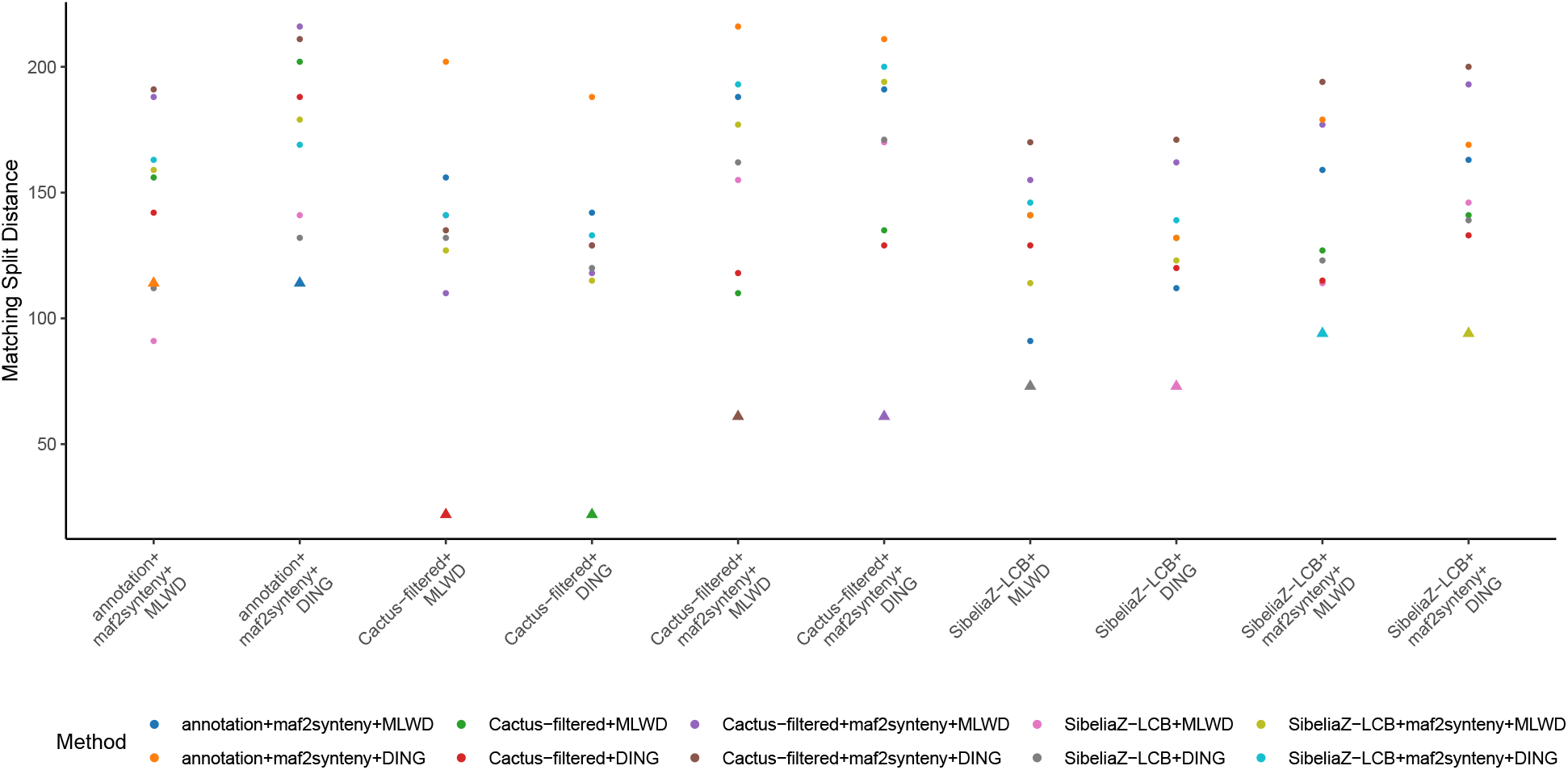
Comparison of fully resolved MLWD and DING trees using Matching Split distance. As in Fig. 5, each point corresponds to the matching split distance between the method indicated at the x position and that denoted by the point’s color. The MS distances between trees built from the same underlying synteny blocks but using a different tree inference method are shown as triangular points. In the majority of cases, the trees closest to each other are in fact these pairs, with the exception of annotation+maf2synteny+MLWD having greater similarity to both trees from SibeliaZ-LCB+maf2synteny than to annotation+maf2synteny+DING.

The only case where the tree inference method mattered more than the block determination method was annotation+maf2synteny+MLWD, where this tree was slightly closer to both SibeliaZ-LCB+MLWD (31% RF, 91 MS) and SibeliaZ-LCB+DING (29% RF, 112 MS) than to its partner tree annotation+maf2synteny+DING (27% RF, 114 MS).

## 4 Discussion

As inferring phylogenies based on large-scale mutations becomes increasingly more feasible in terms of data availability, many questions about the best practices remain unanswered. Before the methods diligently developed by the research community are more broadly adopted, we need more empirical analyses that guide the practitioners in building robust and reliable analysis pipelines. In this paper, we took a step in that direction. Using a dataset of high quality Mtb genomes, we interrogated robustness of methods used for preparing the syntenic blocks, which form the inputs to methods that infer phylogenies using rearrangements.

### 4.1 Limitations of the study and future work

The space of possible methods for obtaining input to rearrangement phylogenies is wide and this study is necessarily incomplete. While we have made an effort to represent the most modern WGA-based strategies for block assignment, many alternatives exist and future work should explore them.

While we considered two sets of parameters for orthology assignment of the annotation blocks, we did not consider alternative assignment methods [Linard et al., 2021]. The full impact of the different methods for annotation, orthology detection, and post-processing of orthology on the eventual phylogenetic inference remains to be explored. Additionally, we limited our study to the use of maf2synteny for the agglomeration of basic homology statements into syntenic blocks. Other methods such as i-ADHoRe3.0 [Proost et al., 2012] and Cyntenator [Rödelsperger and Dieterich, 2010] exist for this purpose. We were unsuccessful in running each of these on our dataset: i-ADHoRe3.0 has an array of parameters and, using the recommended values, we received empty output. Turning to Cyntenator, running default parameters produced only 2 synteny blocks. After adjusting parameters and rerunning, Cyntenator did not complete after a week of running.

Cactus’ requirement of a guide tree to determine the order of pairwise alignment became an important variable for the subsequent inference of the tree from the alignment. It should also be kept in mind that our dataset consists of many isolates from a single species. The yet-unpublished Cactus Pangenome Pipeline is presented as a solution to the problem of guide trees, though rather than a guide tree, it requires specifying a reference genome. This option should be explored as an alternative solution.

The extent to which we could make comparisons to the distance-based trees was limited since we did not have bootstrapping available to be able to identify more strongly supported branches and assess whether concordance improved among them. As a method for bootstrapping distance-based trees exists [Lin et al., 2012b], implementation and application of this method is important to learning more about these trees’ reliability.

To combine methods inferred from two types of data, we relied on the fact that they were fully compatible for highly supported branches. Such simple supertree methods are perfect for interpretation, as one knows each branch has full support from at least one data source and no strong conflict from the other. Nevertheless, full compatibility will not always be achieved, necessitating more complex procedures for obtaining supertree methods [Bininda-Emonds, 2004], which may reduce interpretability. An arguably more principled approach for combining signal is to combine the data and infer one tree from the entirety of the data. This combination, however, is not trivial. We can perhaps concatenate data and use data partitioning, but proper weighting of data is not obvious. Such approaches need to be further explored and compared to the supertree method.

Finally, method evaluation on real data, while free of concern about realism of the data, is complicated by lack of access to the ground truth. Realistic genome simulation remains challenging; options that do exist tend to have important shortcomings such as lack of support, bugs, and limited features. We require a method that supports genome-scale rearrangement as well as gene-scale substitutions, a model for intergenic nucleotide evolution, and the specification of a real genome at the root of the simulated species tree. As far as we know, Evolver [Edgar et al.] is the only option that has anywhere near this feature set, but the software is complex to set up, was designed specifically for eukaryotic genomes, and is no longer supported by its authors. Once improved methods for genome simulation are available, repeating our analyses will allow a more direct measurement of accuracy, at every step of the pipeline.

### 4.2 Practical Lessons

Our results both create cause for caution and room for optimism. It would be ideal if the inferred phylogenies were robust to the method of obtaining blocks and the method of inferring the phylogeny. Instead, we saw that not only the choice of the block assignment method matters, but also some parameter settings of methods can impact the results. The block composition seems sensitive to the parameters of alignment, on one hand, and parameters of the methods used to group alignment segments into larger groups (e.g., maf2synteny), on the other. The block properties, in turn, impact the phylogeny. Furthermore, some parameters, such as the guide tree used for WGA, do not impact block composition in obvious ways but impact the final tree in more subtle ways. Thus, practitioners are encouraged to remain cautious about these choices, and our results call for more extensive empirical analyses.

The apparent importance of the choice of synteny block construction method on the downstream phylogenetic inference should not surprise us. Traditional phylogenetics has long wrestled with impacts of incorrect homology detection and alignment on tree inference [Ogdenw and Rosenberg, 2006, Lunter et al., 2008, Liu et al., 2009, Li-San Wang et al., 2011, Philippe et al., 2017, Springer and Gatesy, 2018], and it would be naive to expect rearrangement phylogenies would be spared that concern. In fact, some of our findings are analogous to similar observations for multiple sequence alignment. For example, the impact of guide trees on final adjacency trees reminds one of the impact of guide trees on multiple sequence alignments and resulting trees [Nelesen et al., 2008, Boyce et al., 2014].

Beyond parameters, our results provide reasons to prefer some methods versus others. Compared to using gene annotations, the alignment-based methods showed superior compatibility with the reference SNP tree and with each other. In contrast, the annotation-based tree has numerous incompatibilities, even among branches with 100% bootstrap support, with the SNP tree. The large number of blocks in the annotation input is cause for concern, and not greatly improved by further aggregation. It appears that annotation pipelines, even run with relaxed settings, fail to make complete orthology calls in our dataset. Such failures to find orthology can easily lead to inconsistent adjacencies across genomes, and perhaps high support for wrong branches. Thus, our results support the use of WGA, in general, and Cactus, in particular, so long as a suitable guide tree can be determined. Moreover, assuming the SNP tree as the reference, our results seem to suggest that maximum-likelihood trees computed with the adjacency-based method MLGO have superior accuracy compared to the distance-based method DING.

Despite the variability that we observed, some encouraging patterns emerged. The Cactus adjacency tree did have high levels of compatibility and a *complementary* signal to the SNP tree, allowing us to combine their highly supported branches into a supertree with more resolution than either tree. It is true that the compatibility is helped by the choice of the SNP tree as the guide, complicating the interpretation of compatibility as accuracy. Nevertheless, if we are not using the SNP tree for benchmarking, this compatibility shows a path forward. Practitioners can infer preliminary substitution-based trees, using reference genomes or even assembly-free methods [Ondov et al., 2016b, Sarmashghi et al., 2019, Lau et al., 2019] and infer a WGA using those as guide tree. The WGA can then be used both with adjacency-based and substitution-based models to infer alternative trees, which when compatible, can easily and unambiguously be combined. When this process is successful, as it was here, the long-standing goal of combining signal from two types of events is achieved.

