## Supplemental Tables and Figures for "Evaluating Impacts of Syntenic Block Detection Strategies on Rearrangement Phylogeny Using M. tuberculosis Isolates"

### Supplementary Material

#### 1 Supplementary Tables and Figures

| Program | Version | Runtime |
|---|---|---|
| Hybran (annotation) | 1.2.0 | 146 minutes |
| Hybran (annotation-relaxed) | | 100 minutes |
| SibeliaZ-LCB | 1.2.2 | 3 minutes |
| Cactus | 1.3.0 | 22 hours 17 minutes |
| DING | 2021-01-26 | 2 hours 17 minutes (SibeliaZ-LCB) - 11 hours, 55 minutes (Cactus(Mash)-filtered) |
| Mash | 2.3 | |
| maf2synteny | 1.0 | |
| MLGO | 1.0 | 1 minute (Cactus(SNP)-filtered+maf2synteny) - 12 minutes (Cactus(SNP)) |
| FastME | 2.1.6.2 | |

Table S1: Versions and running time for software used in this study

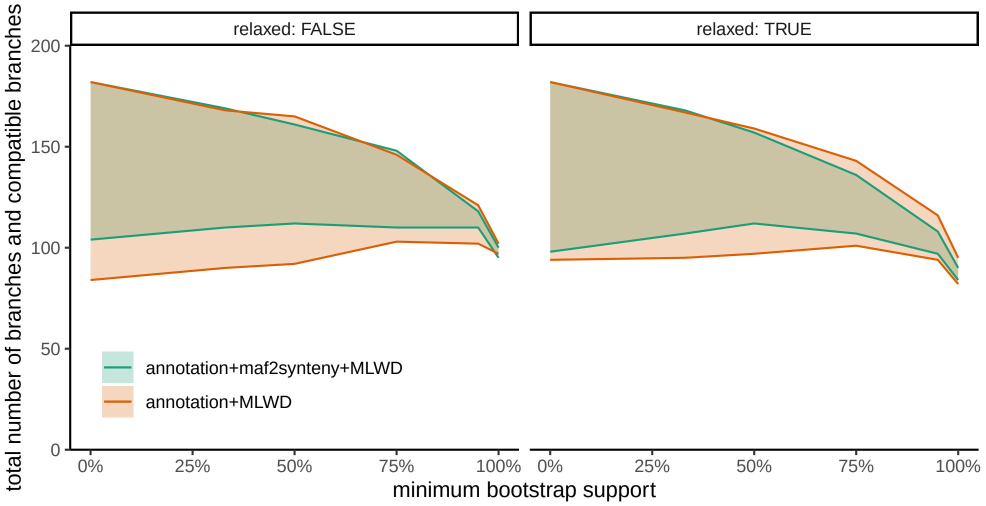


Figure S1: Compatibility of the annotation adjacency trees to the reference SNP tree based on default orthology-mapping parameters (minimum identity 95%, minimum alignment coverage 95%) and relaxed (minimum identity 75%, minimum alignment coverage 66%).

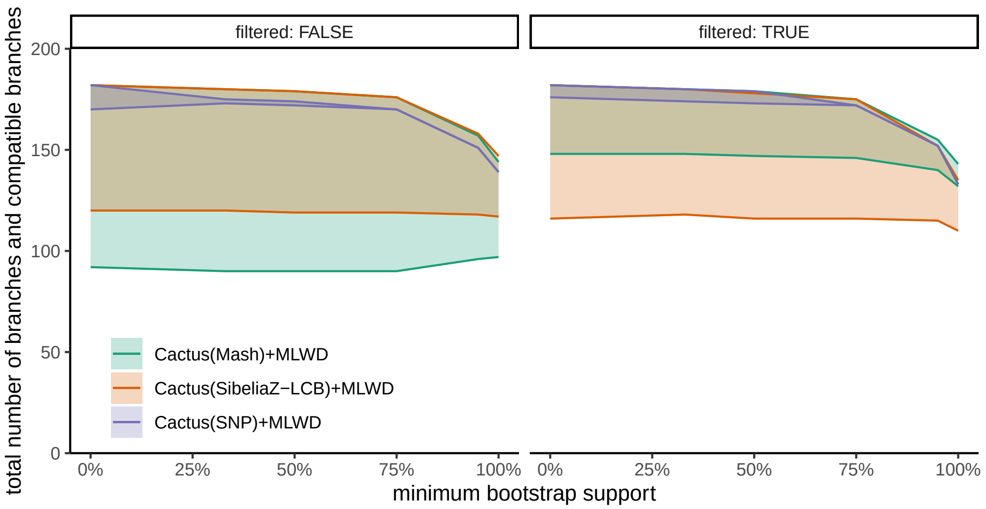


Figure S2: Compatibility of Cactus (with alignment guide tree indicated in parentheses) adjacency trees to the reference SNP tree before and after excluding blocks with fewer than 50 sites. Filtering of small blocks introduces a substantial improvement for Cactus(Mash) and more minimal differences for the alignments based on the other two guide trees.

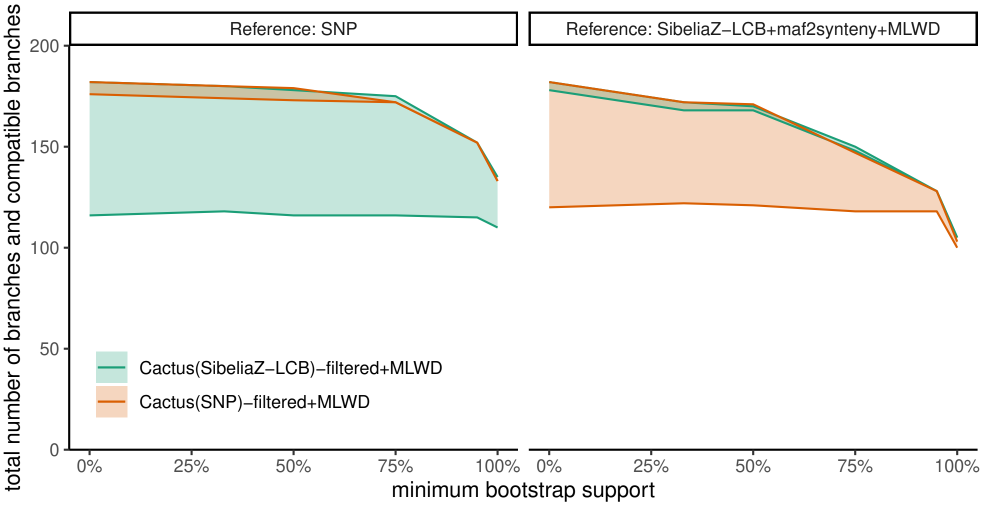


Figure S3: Compatibility of Cactus adjacency trees with different reference trees. Adjacency trees of Cactus alignment exhibit striking degrees of compatibility to the guide tree that was used to create the alignment and much less so to an alternative reference tree, even for branches with high BS.

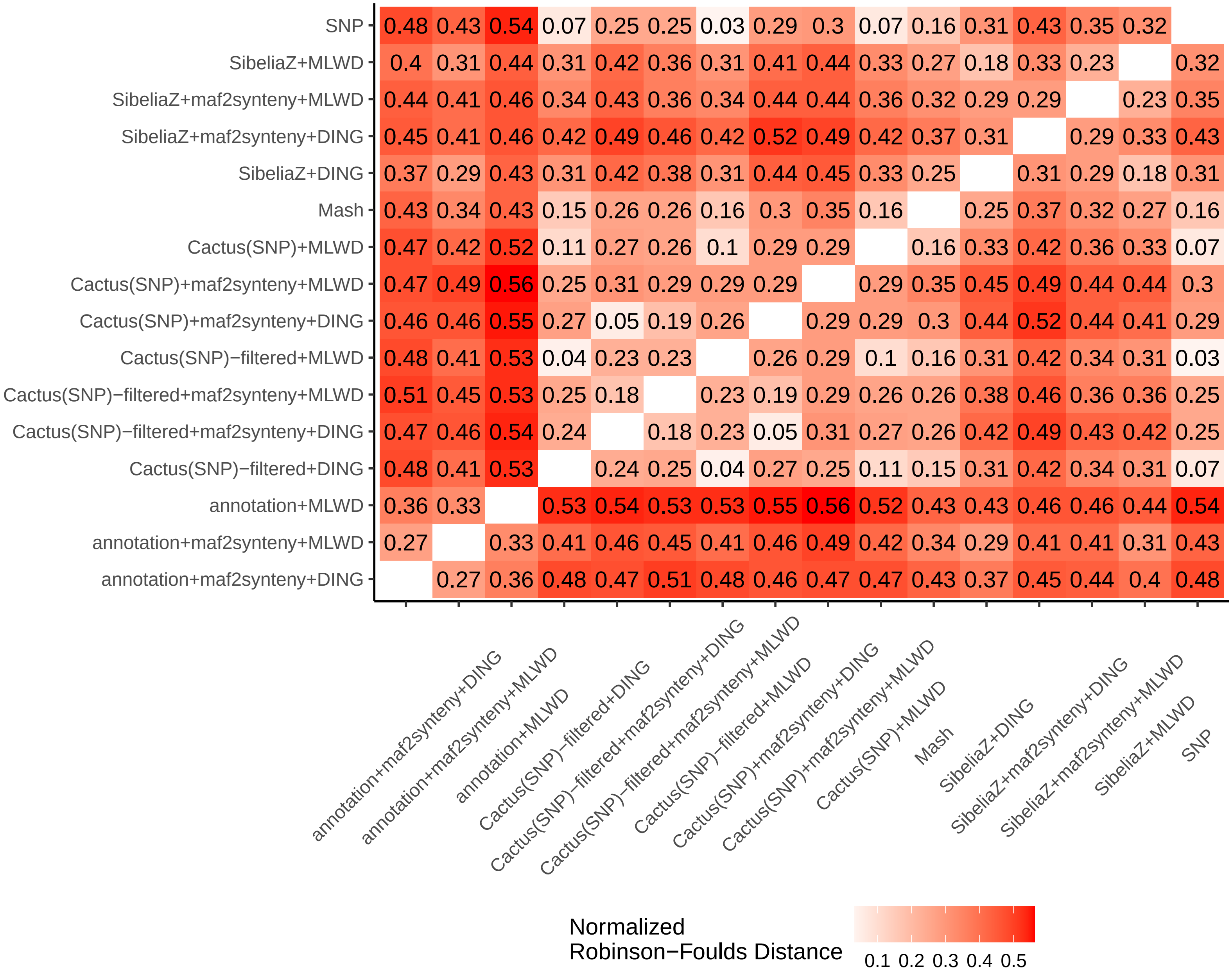


Figure S4: Normalized Robinson-Foulds distances

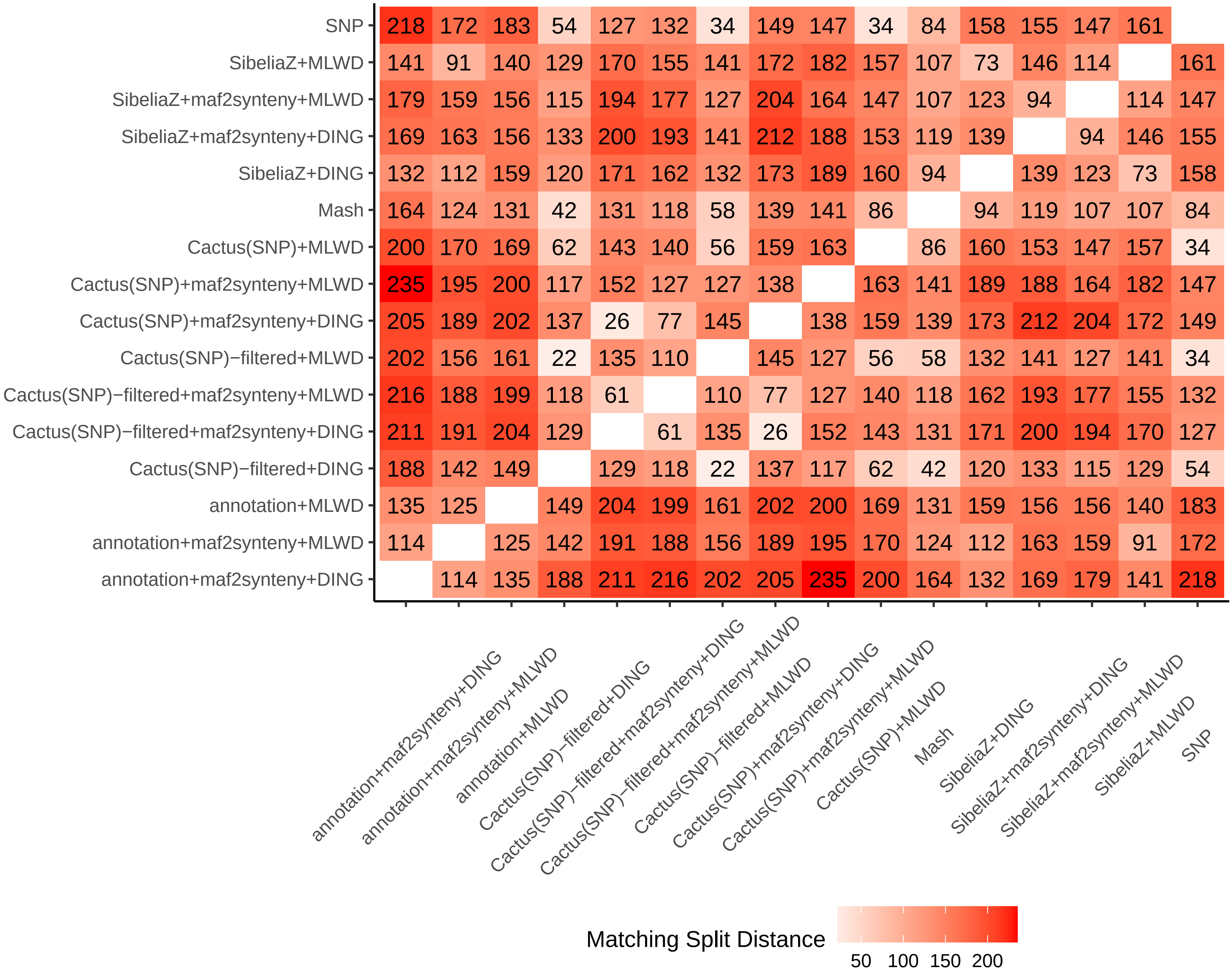


Figure S5: Matching split distances
